# Effect of ligands on stability of H-Ras GTPase

**DOI:** 10.1101/179283

**Authors:** Elizaveta A Kovrigina, Casey O’Connor, Evgenii L Kovrigin

## Abstract

The G domain of a small monomeric GTPase Ras contains a nucleotide-binding pocket and a magnesium-binding site essential for the Ras function in cellular signaling. The G domain also has another (allosteric) ion-binding site on the rear surface of the G domain, which function is still unknown. In this paper, we detailed the effect of calcium and magnesium ions on stability of Ras bound to GDP, GTP, and GTP-mimic GppNHp. We revealed that the remote allosteric ion-binding site contributes very significantly to stability of Ras in the GDP-bound conformation, but nearly not at all—when Ras is bound to a GTP mimic. These findings highlight that further studies of the remote ion-binding site are warranted to reveal its role in the Ras function.

---

Ras is a small monomeric GTPase involved in cellular signaling of growth, proliferation, and differentiation^*1*^. Ras is composed of the 20 kDa cytosolic domain, G domain, and the membrane-bound lipidated C-terminal peptide of 22-23 aminoacids^*2-4*^. The G domain (depicted in Figure 1) functions as a molecular switch, which has high affinity to its binding partners (effectors) in the active, GTP-bound conformation, and loses its affinity in the inactive, GDP-bound state^*4*^.

**Figure 1.**
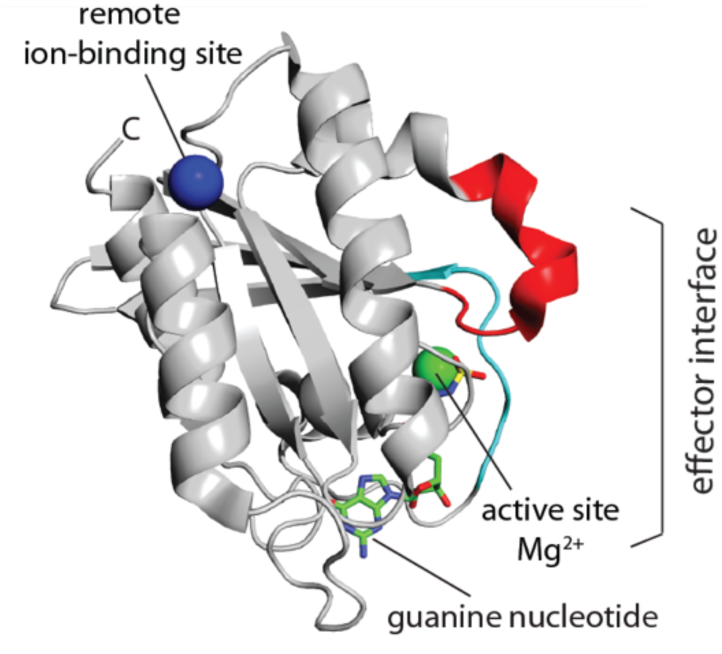
Molecular model of the G-domain of H-Ras GTPase (residues 1-166) in complex with GppNHp, Mg^2+^, and Ca^2+^ (PDBID 3K8Y).

The active conformational state of the G domain bound to GTP is transient because of the intrinsic GTPase activity^*5, 6*^, which is stimulated by the activator protein, GAP, in a cellular context^*7*^. G domain is slow to spontaneously exchange the hydrolysis reaction product, GDP, even if GTP is abundant in the cytosol; the GDP release is accelerated by another catalyst in the cell—guanine nucleotide exchange factor, GEF^*7*^. Along with the nucleotide, the G domain binds a magnesium ion at the active site (green sphere in Figure 1), which significantly increases guanine nucleotide affinity and is critical for GTPase function^*5*^. Recently, one more binding site for the divalent ions was identified in the region of the Ras macromolecule opposite to the effector-interaction interface^*8, 9*^ (remote ion-binding site; blue sphere in Figure 1). We and others demonstrated that binding of ions at this remote site is allosterically coupled to the GTPase pocket^*8, 9*^. In particular, the H isoform of GDP-bound Ras had 2.5x stronger affinity for divalent ions at this site than the GTP-mimicked form, H-Ras-GppNHp^*8*^.

Ligand binding has long been known as a crucial factor defining stability of Ras structure but detailed studies were limited to the effect of GDP and magnesium^*10*^. In this report, we tested stability of the H-Ras G domain, residues 1-166, (in the following text—Ras) in the presence and absence of mono- and divalent ions, guanine nucleotides GDP and GTP, as well as the slowly hydrolysable GTP mimic, GppNHp.

## Results and Discussion

The golden standard in analysis of protein-ligand interactions is the isothermal titration calorimetry, ITC^*11*^. However, this technique is quite slow and requires relatively large samples. The fundamental linkage between stability of the protein structure in a complex with the ligand and affinity of the ligand interaction^*12, 13*^ allows for detection of ligand binding through analysis of relative stability of different protein-ligand complexes. The central idea of this method is that the higher binding affinity (at the room temperature) translates into the higher temperature of protein unfolding because interactions with the ligand make the protein structure more stable. Since protein unfolding may be sensitively followed by fluorescence, this approach could use very small protein samples and become very scalable if implemented with temperature-controlled fluorescent plate-readers. One simple yet indirect way to detect protein unfolding relies on a strong increase in the fluorescence yield of the hydrophobic fluorescent dyes upon their binding to exposed hydrophobic residues of the unfolded polypeptides. The commercial protein dye CYPRO Orange is most commonly utilized because its absorbance and emission bands are in the visible spectral range—suitable for use in the commercial real-time PCR instruments, which are temperature-controlled multi-well fluorometers by design.

The detection of protein-ligand interactions through protein thermal unfolding detected with CYPRO Orange in the real-time PCR machines was first introduced to optimize conditions for X-ray crystallography, and—for ligand screening in drug design (termed “differential scanning fluorometry”, DSF, or the “thermal shift assay”)^*14-16*^. However, since the unfolding transition in the presence of CYPRO is completely irreversible, the midpoint of the “unfolding” part of the DSF profile may only serve as an estimate of the temperature at which ΔG(F->U)=0, and the shape of the transition cannot be expected to conform to the equations for the folding-unfolding equilibrium. Therefore, the DSF data is non-equilibrium and thus cannot be subject to a rigorous thermodynamic linkage analysis. With this limitation in mind, we will use the temperature of middle of the rising part a DSF profile (calling it “unfolding midpoint”) as a gauge of the relative affinity of Ras-ligand interactions.

### Nucleotides and magnesium

Since it is difficult to prepare nucleotide and magnesium-free Ras due to its instability, we started with the 10 μM Ras-GDP that was treated with 1 mM EDTA to completely remove magnesium complexed with the protein (in this sample, the GDP is present in an equimolar ratio to Ras). Black trace in Figure 2 indicates that stability of the magnesium-free Ras-GDP is relatively low with the unfolding midpoint of only 44°C (Table 1). Addition of excess GDP to 1 mM in solution dramatically stabilizes Ras (by 16 °C, blue trace). When magnesium chloride is added to 6 mM in the next step, the free concentration of magnesium ion becomes approximately 5 mM (because 1 mM is sequestered by 1 mM EDTA). Cyan trace in Figure 2 reveals further dramatic stabilization of Ras in these conditions. The total effect of two cognate ligands on the unfolding midpoint is nearly 25°C, which confirms intimate involvement of the guanine nucleotide and magnesium in maintenance of the native Ras conformation, as reported by Zhang and Matthews^*10*^.

**Figure 2.**
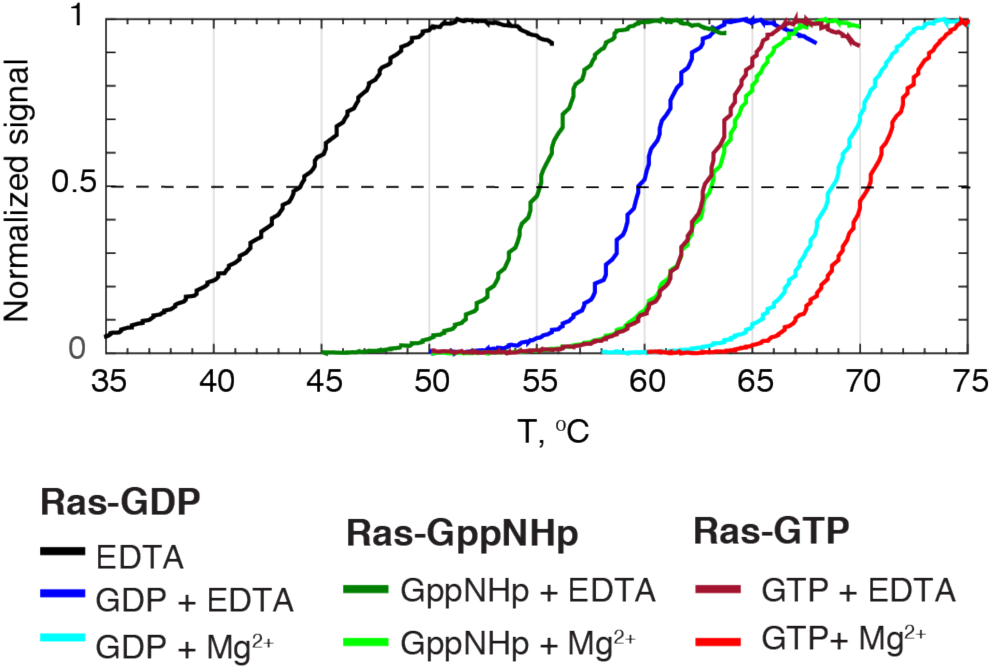
DSF profiles of Ras in the presence of different guanine nucleotides and variable concentrations of magnesium. Pre- and post-transition regions of the DSF curves were clipped for visual clarity; the resulting profiles were normalized to a unity amplitude.

**Table 1.**
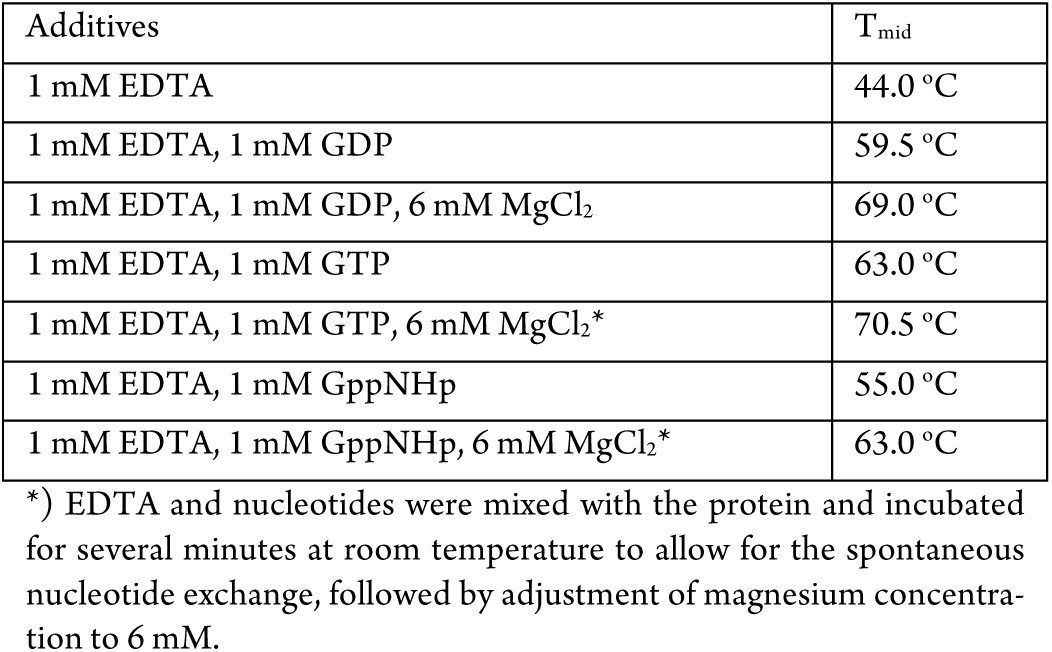
Unfolding midpoints for Ras in the absence and presence of guanine nucleotides and magnesium. Midpoint determination accuracy: 0.5 ° C.

| Additives | T<sub>mid</sub> |
| --- | --- |
| 1 mM EDTA | 44.0 °C |
| 1 mM EDTA, 1 mM GDP | 59.5 °C |
| 1 mM EDTA, 1 mM GDP, 6 mM MgCl<sub>2</sub> | 69.0 °C |
| 1 mM EDTA, 1 mM GTP | 63.0 °C |
| 1 mM EDTA, 1 mM GTP, 6 mM MgCl<sub>2</sub>* | 70.5 °C |
| 1 mM EDTA, 1 mM GppNHp | 55.0 °C |
| 1 mM EDTA, 1 mM GppNHp, 6 mM MgCl<sub>2</sub>* | 63.0 °C |

\*) EDTA and nucleotides were mixed with the protein and incubated for several minutes at room temperature to allow for the spontaneous nucleotide exchange, followed by adjustment of magnesium concentration to 6 mM.

To assess stability of the GTP-bound Ras one must consider that GTP is a natural substrate of Ras GTPase enzyme, therefore is highly instable in the complex with Ras^*5, 6, 17*^. To shorten the time between Ras-GTP sample preparation and measurement, we performed EDTA-assisted nucleotide exchange^*18*^ directly in the well of the optical plate. This method is based on ability of EDTA to remove magnesium from the nucleotide-binding pocket, which accelerates nucleotide dissociation. Therefore, in the presence of the excess GTP in solution, the GDP in Ras is rapidly exchanged by the mass action. The following addition of magnesium “locks” the GTP in the binding site by increasing nucleotide-binding affinity of Ras (as well as activating GTPase function)^*5*^. The dark red trace in Figure 2 shows that the Ras-GTP complex is a very stable species, which is further stabilized when complexed with magnesium (bright red trace).

To characterize the effect of a slowly hydrolysable GTP mimic, GppNHp, on stability of Ras, we performed a similar exchange of endogenous GDP with the excess GppNHp. The geometry and metal-coordination properties of the β, γ-imidotriphosphate chain in GppNHp is different from the triphosphate chain in GTP, which is one reason for slow hydrolysis of GppNHp. This difference also manifests itself in a relatively small degree of stabilization of protein structure both by nucleotide analog itself as well as in the presence of magnesium (dark green and green traces in Figure 2). The relative stabilizing effects of GppNHp, GDP, and GTP (summarized in Table 1) are in agreement with relationship of their affinities for Ras as reported by Scherer et al.^*17*^.

### Interaction with the Ras-binding domain

To demonstrate that Ras protein exchanged with GTP or GppNHp in our samples is functional, we tested its interaction with the cognate effector domain, Ras-binding domain of c-Raf-1, RBD. Figure 3 illustrates the effect of Ras-binding domain interaction on Ras stability. The DSF profile for RBD did not reveal any transitions as probed by CYPRO Orange binding (Supporting Figure 1), likely, because RBD is a fragment of a larger protein and does not have a hydrophobic core well protected from CYPRO in the native state. Therefore, the observed transition on the DSF profile of the Ras-RBD mixture is mostly contributed by Ras unfolding. As anticipated for the Ras-GDP sample, there is little effect of RBD addition as the GDP-bound conformation has low affinity for RBD (black and green traces in Figure 3). In contrast, the Ras-GppNHp is stabilized by nearly 20°C, which is expected for the high-affinity complex^*19*^ (blue and red traces).

**Figure 3.**
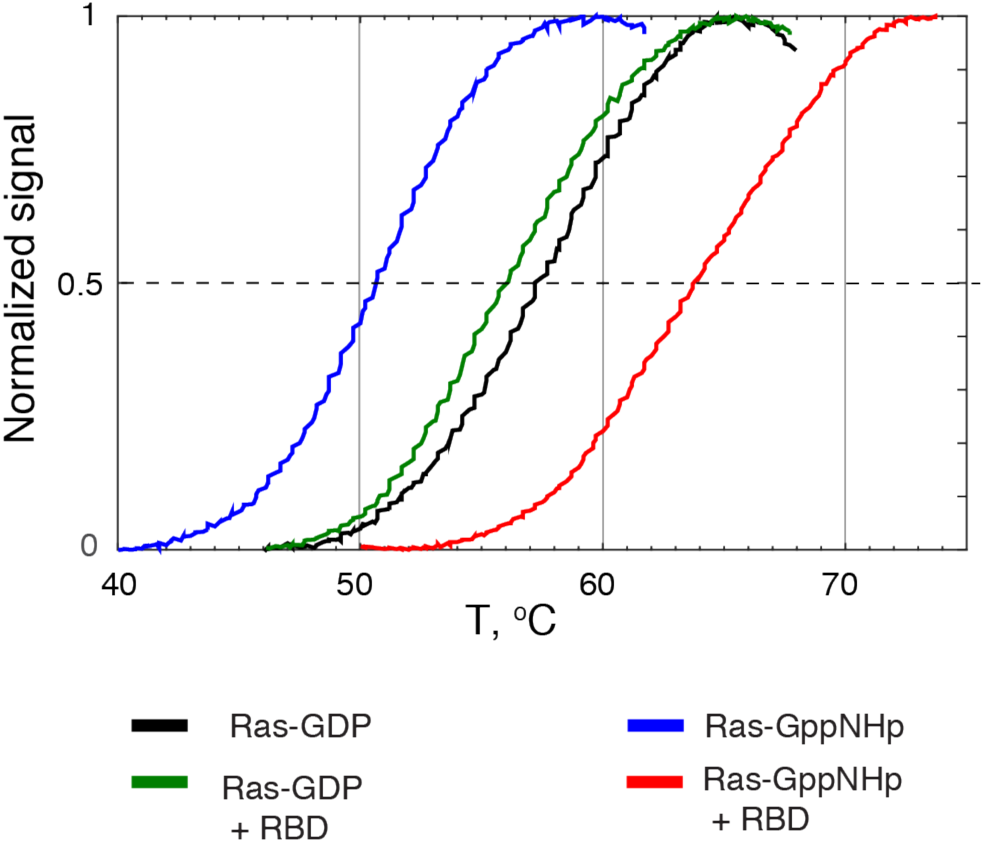
DSF melting profile for Ras in presence of Raf-RBD in the presence of 2 mM MgCl_2_.

A large increase in the transition midpoint temperature observed for Ras-GppNHp-RBD complex is, likely, a combined effect of stabilization due to the high affinity interaction as well as the resulting slow unfolding kinetics. Slow dissociation of the complex may leads to a scan-rate-dependent shift of the profile to the higher temperatures because protein molecules may not have enough time to unfold^*20, 21*^. Therefore, the transition midpoint of the DSF profiles should be considered as an *upper estimate* for the unfolding temperature.

### Ions

To characterize overall ionic preferences the G domain in H-Ras, we explored its stability in a range of inorganic salt solutions with different combinations of anions and cations. Figure 4 demonstrates stability of Ras complexed with GDP (top panel) and the GTP mimic, GppNHp (bottom panel). Addition of magnesium ion significantly stabilizes both forms of Ras (black for 0 mM and red/orange — for 25 mM magnesium acetate/chloride, accordingly). Calcium ions stabilize Ras-GDP while it has small destabilizing effect on Ras-GppNHp. We previously reported similar binding affinities of the remote ion-binding site for calcium and magnesium in both GDP and GppNHp forms^*8*^. Therefore, the observed differential effect calcium on Ras-GDP and Ras-GppNHp must be attributed to the Ca^2+^ binding at the GTPase pocket. The negatively charged counter-ions (chloride or acetate) as well as monovalent sodium do not affect stability of either Ras form, which is in agreement with our earlier report^*8*^.

**Figure 4.**
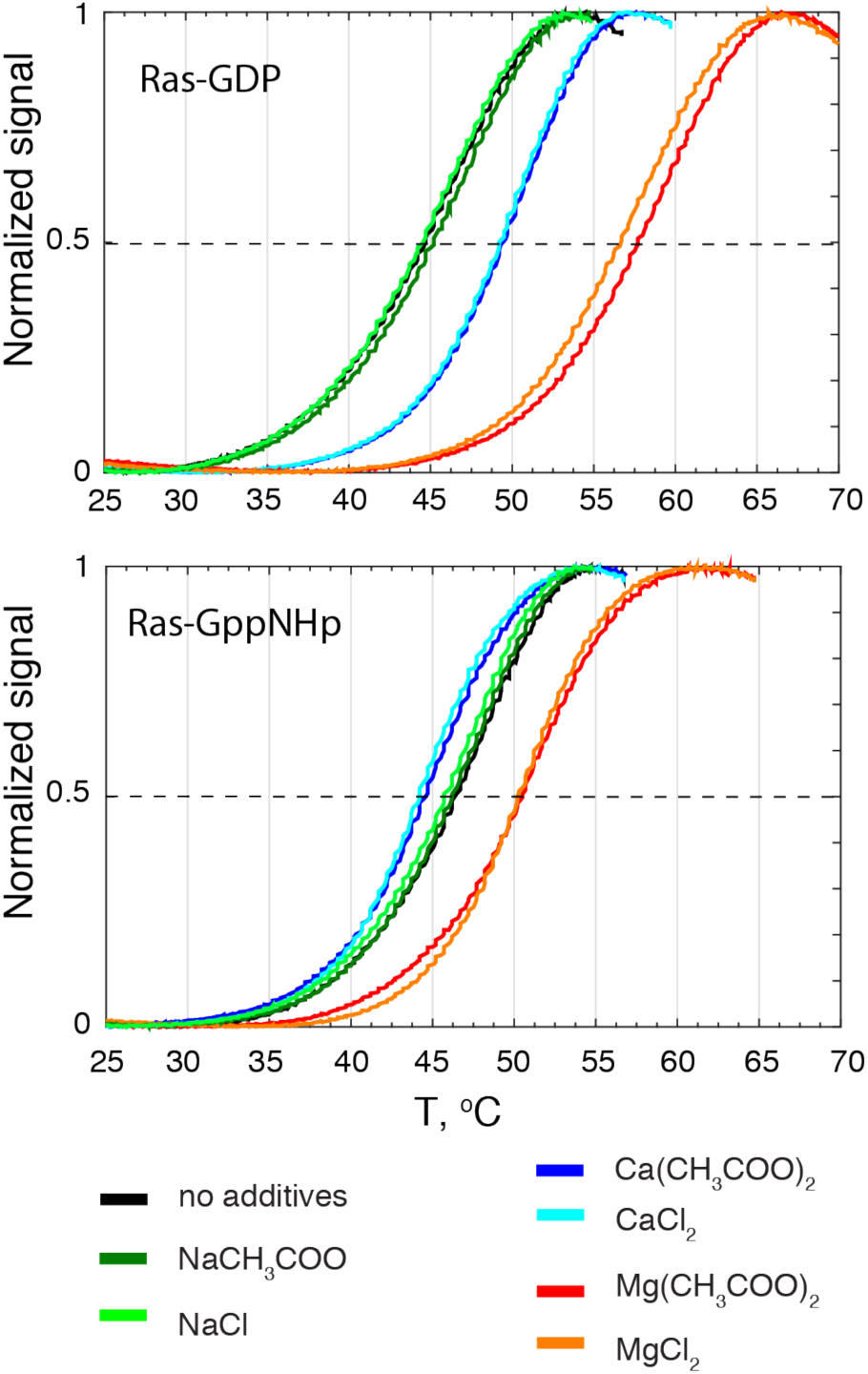
DSF profiles of the Ras-GDP(**Top)** and Ras-GppNHp(**Bottom**) in the 50 mM Hepes pH 7.2 buffer (black) in the presence of 25 mM calcium acetate (blue), CaCl_2_ (cyan), magnesium acetate (red), magnesium chloride (orange), sodium acetate (dark green), and sodium chloride (bright green). Black traces are over-lapped with green traces in both graphs.

### Titration with magnesium

The two ion-binding sites of G domain were reported to have different affinities for a magnesium ion: the micromolar K_d_ at the GTPase pocket^*6*^ and millimolar—at the remote site^*8*^. To further explore interplay of the ionic interactions at these two sites and their contribution to a structural stability of the G domain, we performed a titration of GDP and GTP-mimicked forms of Ras with MgCl_2_. For the titration, we prepared independent solutions for all titration points in triplicate on one 96-well plate and measured stability of all samples simultaneously in one temperature scan.

Figure 5 shows the dependence of the unfolding temperature on the concentration of magnesium ion in solutions. Ras-GppNHp (red circles) shows the stabilizing effect of magnesium that plateaus after 100 μM of MgCl_2_. The magnesium ion at the GTPase site is bound with a micromolar dissociation constant (2.8 μM for Ras-GDP)^*6*^, which is close to a midpoint of the Ras-GppNHp stabilization curve. Thus, as we gradually raise the MgCl_2_ concentration to a high-micromolar level, the protein stability increases until the ion-binding site at the GTPase pocket becomes saturated. Once the site is filled, the further increase of magnesium concentration (above 0.1 mM) does not affect stability of the GTP-mimicked Ras conformation.

**Figure 5.**
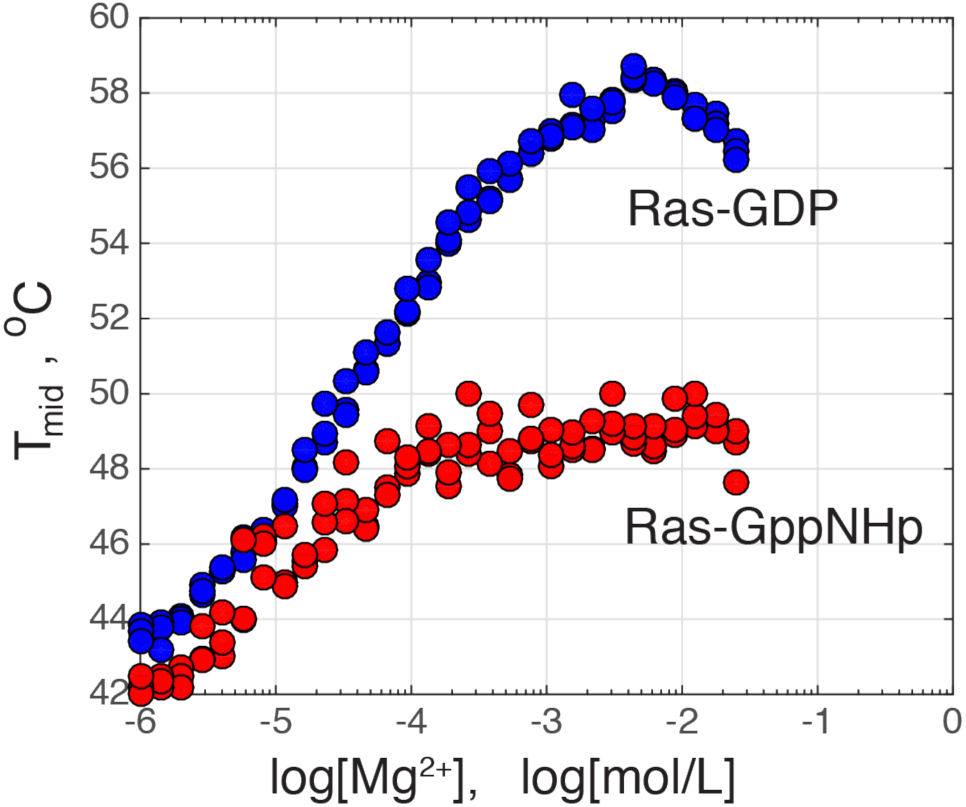
Unfolding midpoint of the Ras G domain as a function of the MgCl_2_ concentration. Circles reflect midpoints of transitions observed in individual wells on the 96-well plate. All conditions were sampled in triplicate. Size of a symbol reflects an average 0.5°C accuracy of the unfolding midpoint values.

The Ras-GDP stabilization curve in Figure 5 (blue circles) reveals a remarkable difference from the Ras-GppNHp profile. While both are similarly stabilized upon saturation of the magnesium site in the GTPase pocket, the Ras-GDP curve stability continues to increase until nearly 10 mM MgCl_2_. We hypothesize that this continuing rise of protein stability is a contribution of the remote ion-binding site, which has millimolar dissociation constant^*8*^. We reported that magnesium ion has 2.5 higher dissociation constant in Ras-GppNHp than in Ras-GDP, which agrees with stronger stabilization of the Ras-GDP by magnesium in Figure 5. These observation highlight a potentially important role of the remote ion-binding site in function of the G domain, which calls for a more rigorous thermodynamic analysis by Differential Scanning Calorimetry^*20, 22, 23*^, ITC combined with site-directed mutagenesis.

## Summary

In this paper, we surveyed stabilization effects of nucleotides and inorganic ions on the G domain of H-Ras. We observed a very distinct impact of the remote binding site on stability of the G domain in complex with the GDP relatively to the GTP-mimic, which may imply an important role for the remote site in Ras structure and function.

## Methods

H-Ras G domain was expressed, isolated and purified following our published protocol^*24*^ and dialized into the magnesium-free working buffer (50 mM HEPES pH 7.2, 1 mM DTT, 0.01% of NaN_3_). For the magnesium titration, the Ras-GppNHp sample was prepared as described^*8*^ and dialyzed to the working buffer. The gene for Ras-binding domain of c-Raf-1, RBD, was obtained from OriGene. The RBD was expressed as the glutathione S-transferase fusion protein, subject to thrombin cleavage, and purified following the published protocols^*19*^. Supporting Figure 2 shows purity of the Ras and RBD preparations.

The DSF unfolding experiments were performed in the Agilent MX3005p QPCR System (Agilent). CYPRO Orange (Sigma S5692) was used in a ratio of 1:1000 by volume. Protein concentration in DSF experiments were 2-10 μM. Scan rate was 1°C/min and the instrument used a standard SYBR Green filter set. The EDTA, nucleotide, and salt stocks were prepared on the working buffer, and their pH value was adjusted to 7.2 prior to experiments.

## Acknowledgement

ELK acknowledges the Regular Research Grant 2012 from Committee on Research (COR), Marquette University.

## Supporting Information

### Supporting Figures

**Supporting Figure 1.**
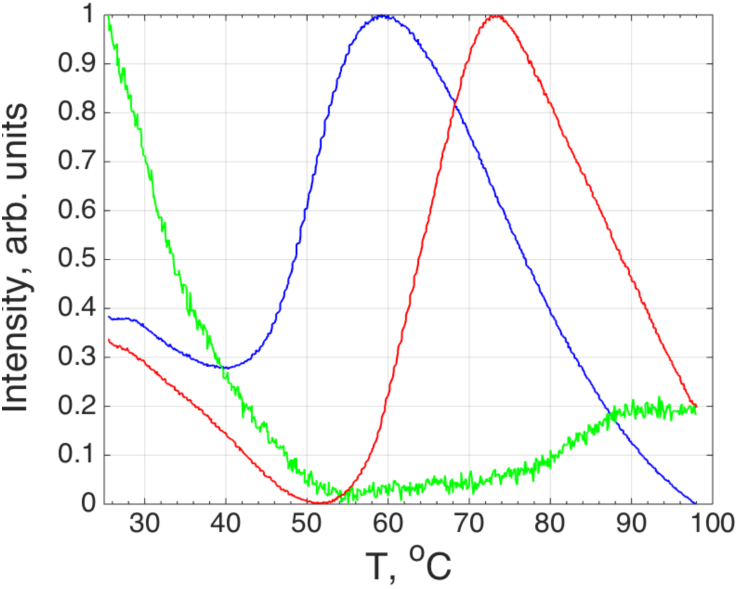
Full (untrimmed) normalized DSF profiles for Ras-GppNHp (blue), c-Raf-1 RBD (green) and Ras-RBD complex (red).

**Supporting Figure 2.**
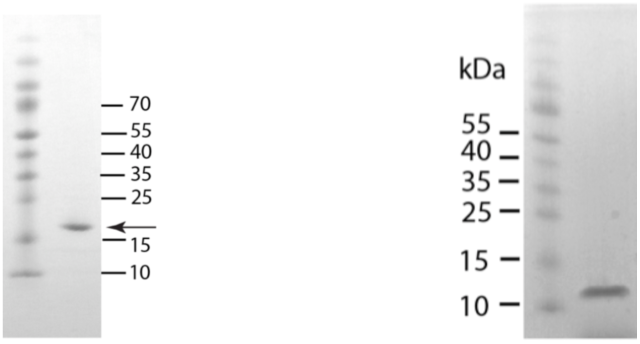
SDS-PAGE of the purified H-Ras G domain preparation (left) and RBD preparation (right) used for binding studies.

## References

[1] Colicelli, J. (2004) Human RAS Superfamily Proteins and Related GTPases, Sci. STKE 2004, re13.

[2] Hancock, J. F., Magee, A. I., Childs, J. E., and Marshall, C. J. (1989) All ras proteins are polyisoprenylated but only some are palmitoylated, Cell 57, 1167–1177.

[3] Eisenberg, S., Laude, A. J., Beckett, A. J., Mageean, C. J., Aran, V., Hernandez-Valladares, M., Henis, Y. I., and Prior, I. A. (2013) The role of palmitoylation in regulating Ras localization and function, Biochemical Society Transactions 41, 79–83.

[4] Wittinghofer, A., and Vetter, I. R. (2011) Structure-Function Relationships of the G Domain, a Canonical Switch Motif, In Annual Review of Biochemistry (Kornberg, R. D., Raetz, C. R. H., Rothman, J. E., and Thorner, J. W., Eds.), pp 943–971, Annual Reviews, Palo Alto.

[5] John, J., Sohmen, R., Feuerstein, J., Linke, R., Wittinghofer, A., and Goody, R. (1990) Kinetics of interaction of nucleotides with nucleotide-free H-ras p21, Biochemistry 29, 6058–6065.

[6] John, J., Rensland, H., Schlichting, I., Vetter, I., Borasio, G. D., Goody, R. S., and Wittinghofer, A. (1993) Kinetic and structural analysis of the Mg(2+)-binding site of the guanine nucleotide-binding protein p21H-ras, The Journal Of Biological Chemistry 268, 923–929.

[7] Bos, J. L., Rehmann, H., and Wittinghofer, A. (2007) GEFs and GAPs: Critical elements in the control of small G proteins, Cell 129, 865–877.

[8] O’Connor, C., and Kovrigin, E. L. (2012) Characterization of the Second Ion-Binding Site in the G Domain of H-Ras, Biochemistry 51, 9638–9646.

[9] Buhrman, G., O’Connor, C., Zerbe, B., Kearney, B. M., Napoleon, R., Kovrigina, E. A., Vajda, S., Kozakov, D., Kovrigin, E. L., and Mattos, C. (2011) Analysis of binding site hot spots on the surface of Ras GTPase, J Mol Biol 413, 773–789.

[10] Zhang, J., and Matthews, C. R. (1998) Ligand Binding Is the Principal Determinant of Stability for the p21H-ras Protein, Biochemistry 37, 14881–14890.

[11] Velazquez-Campoy, A., Ohtaka, H., Nezami, A., Muzammil, S., and Freire, E. (2004) Isothermal titration calorimetry, Curr Protoc Cell Biol Chapter 17, Unit 17 18.

[12] Luque, I., and Freire, E. (2000) Structural stability of binding sites: Consequences for binding affinity and allosteric effects, Proteins-Structure Function and Genetics, 63–71.

[13] Freire, E. (1998) Statistical thermodynamic linkage between conformational and binding equilibria, In Advances in Protein Chemistry, Vol 51, pp 255–279.

[14] Niesen, F. H., Berglund, H., and Vedadi, M. (2007) The use of differential scanning fluorimetry to detect ligand interactions that promote protein stability, Nat. Protoc. 2, 2212–2221.

[15] Vedadi, M., Niesen, F. H., Allali-Hassani, A., Fedorov, O. Y., Finerty, P. J., Wasney, G. A., Yeung, R., Arrowsmith, C., Ball, L. J., Berglund, H., Hui, R., Marsden, B. D., Nordlund, P., Sundstrom, M., Weigelt, J., and Edwards, A. M. (2006) Chemical screening methods to identify ligands that promote protein stability, protein crystallization, and structure determination, Proceedings of the National Academy of Sciences of the United States of America 103, 15835–15840.

[16] Huynh, K., and Partch, C. L. (2015) Analysis of protein stability and ligand interactions by thermal shift assay, Curr Protoc Protein Sci 79, 28 29 21–14.

[17] Scherer, A., John, J., Linke, R., Goody, R. S., Wittinghofer, A., Pai, E. F., and Holmes, K. C. (1989) Crystallization and Preliminary X-ray Analysis of the Human c-H-ras-Oncogene Product p21 Complexed with GTP Analogues, Journal of Molecular Biology 206, 257–259.

[18] Hall, A., and Self, A. J. (1986) The effect of Mg2+ on the guanine nucleotide exchange rate of p21N-ras, J. Biol. Chem. 261, 10963–10965.

[19] Herrmann, C., Martin, G. A., and Wittinghofer, A. (1995) Quantitative Analysis of the Complex between p21 and the Ras-binding Domain of the Human Raf-1 Protein Kinase, J. Biol. Chem. 270, 2901–2905.

[20] Potekhin, S. A., and Kovrigin, E. L. (1998) Folding under inequilibrium conditions as a possible reason for partial irreversibility of heat-denatured proteins: computer simulation study, Biophysical Chemistry 73, 241–248.

[21] Potekhin, S. A., and Kovrigin, E. L. (1998) Influence of kinetic factors on heat denaturation and renaturation of biopolymers, Biofizika 43, 223–232.

[22] Privalov, P. L. (2009) Microcalorimetry of proteins and their complexes, Methods Mol Biol 490, 1–39.

[23] Privalov, P. L., and Potekhin, S. A. (1986) Scanning microcalorimetry in studying temperature-induced changes in proteins, Methods Enzymol 131, 4–51.

[24] O’Connor, C., and Kovrigin, E. L. (2008) Global conformational dynamics in Ras, Biochemistry 47, 10244–10246.

